# RfaA (YqhY), a novel adaptor protein, controls metabolite-sensitive protein degradation in *Bacillus subtilis*

**DOI:** 10.1101/2024.02.19.580884

**Authors:** Dennis Wicke, Sabine Lentes, Cornelia Herrfurth, Dominik Tödter, Lorena Stannek, Fabian M. Commichau, Ivo Feussner, Jörg Stülke

## Abstract

The carboxylation of acetyl-CoA is the committed step of fatty acid synthesis catalyzed by the multisubunit acetyl-CoA carboxylase (ACCase). However, the mechanisms that control the activity of this enzyme are poorly understood. Here, we identify the so far unknown protein RfaA (YqhY) of the model bacterium *Bacillus subtilis* as a regulator of fatty acid <u>a</u>cquisition that targets the AccC subunit of ACCase for degradation. RfaA interacts with both AccC and the ClpE unfoldase subunit of the ClpEP protease complex. While the former interaction is not sensitive to the physiological conditions, RfaA interacts with ClpE only in the absence of the global amino group donor glutamate. This results in the degradation of AccC in the absence of glutamate. The inactivation of the *rfaA* gene results in the accumulation of fatty acids in the cell and in the formation of lipid droplets which are toxic for the bacteria. The reduced fatty acid synthesis in the absence of glutamate as a result of RfaA-and ClpEP-dependent degradation of AccC thus prevents intoxication of the cells by fatty acids. Our findings suggest that the degradation of enzymes that catalyze the committed step of biosynthetic pathways might be an important mechanism to control cellular homeostasis.

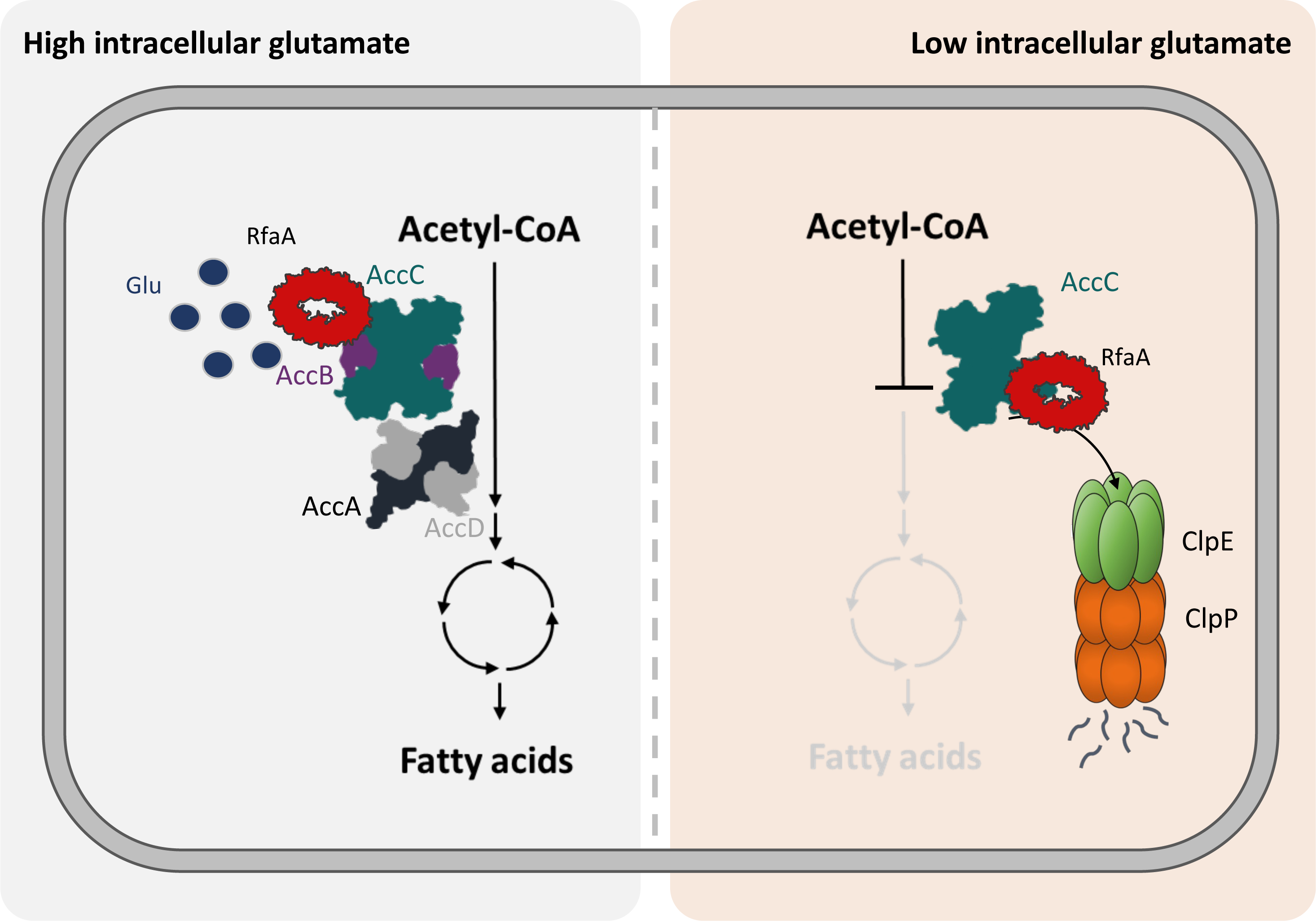

## Introduction

Even after decades of intensive research, the function of a large number of proteins remains unknown. This is even the case for well-studied model microorganisms such as *Escherichia coli*, *Bacillus subtilis*, and the bakers yeast *Saccharomyces cerevisiae* ^1^ as well as for the artificial minimal organism *Mycoplasma mycoides* JCVI-syn3A ^2^. For the Gram-positive model bacterium *B. subtilis*, the function of about 25% of all proteins is completely unknown or only very vaguely understood ^3^. Many of these proteins are only very poorly expressed suggesting that they play a role only under very specific conditions ^4^. However, several of the unknown proteins are highly expressed under a wide range of conditions indicating that these proteins are of vital importance for the physiology of *B. subtilis* ^5,6,7^. The assignment of functions to unknown proteins is often difficult and is substantially supported by the availability of some initial information that guides the direction of further research ^8^.

The so far unknown *B. subtilis* protein YqhY is encoded by one of the most strongly expressed genes in this bacterium suggesting that it plays an important role in the cell. Orthologs of YqhY are present in essentially all species of the Firmicutes and in many other bacteria suggesting that these proteins share a conserved function. This protein family has been named Asp23 protein family based on the *Staphylococcus aureus* Asp23 protein that is induced upon alkaline shock ^9^. The *S. aureus* Asp23 protein has been implicated in cell envelope homeostasis; however, no specific function or activity has been identified ^9^. Interestingly, the bacteria that encode Asp23 family proteins, typically possess two, sometimes even more paralogs of the protein ^10^. *B. subtilis* encodes two proteins of the Asp23 family, YqhY and YloU that share 32% amino acid identity. The two proteins are encoded in conserved operons with the *accBC* genes and the *fakA* gene, respectively. The *accBC* genes encode subunits of the essential acetyl-CoA carboxylase (ACCase), whereas *fakA* encodes a fatty acid kinase that allows the utilization of fatty acids from the environment for lipid biosynthesis ^11,12^. This conserved gene clustering suggests a functional link between the Asp23 proteins and the two distinct mechanisms for fatty acid acquisition for phospholipid biosynthesis, *i. e. de novo* biosynthesis or the use of exogenous fatty acids. Both *B. subtilis* genes that encode proteins of the Asp23 family are strongly expressed suggesting that they are of vital importance for the cell. The *yqhY* gene belongs even to the group of the most strongly expressed genes of unknown function that were identified as important targets for functional analysis ^7^. The idea that YqhY is likely very important for *B. subtilis* is underlined by the observation that the bacteria readily acquire suppressor mutations upon loss of the YqhY protein. The analysis of a comprehensive set of such mutants revealed mutations affecting all subunits of ACCase, which catalyzes the initial step in fatty acid biosynthesis ^13^. This observation further strengthens the idea of a possible link between YqhY and lipid synthesis. The conserved *accBC-yqhY* operon organization as well as the appearance of suppressor mutations affecting the ACCase are the initial information that can provide a primer for identification of the molecular function of YqhY.

All living cells are separated by a cell membrane from their environment. In most cases, the membranes consist of phospholipids with fatty acids as a major building block and integrated proteins. To assure integrity of the cell, the synthesis of the cell envelope has to be tightly controlled and needs to be coupled to the supply of building blocks from central metabolism ^14,15,16^. This is easily achieved with respect to the availability of carbon sources as acetyl-CoA and glycerol-3-phosphate both derived from glycolysis are the major precursors for lipid biosynthesis. On the other hand, essentially nothing is known about the impact of other macroelements such as nitrogen, phosphorus, or sulphur on lipid biosynthesis in bacteria. These macroelements also need to be available for cell growth and no lipid biosynthesis is required especially if nitrogen is limiting. In *B. subtilis* and related Gram-positive bacteria, fatty acid biosynthesis is controlled the transcription repressor FapR, that prevents expression of fatty acid biosynthetic genes if the first dedicated intermediate of the pathway, malonyl-CoA, is limiting. In contrast, the genes of the FapR regulon are induced upon malonyl-CoA biosynthesis ^17^. However, the regulation of malonyl-CoA synthesis by the essential ACCase has so far not been studied.

In this study, we have identified the availability of glutamate as the factor that determines whether the cells require YqhY or not. In the absence of glutamate, the loss of YqhY results in the accumulation of large amount of lipids that poison the cell. YqhY helps to prevent this intoxication by fatty acids in the absence of glutamate by triggering the degradation of the biotin carboxylase subunit (AccC) of the ACCase complex. YqhY acts as an adaptor protein that delivers the AccC protein in a glutamate-dependent manner for degradation to the ClpEP protease complex.

## Results

### Glutamate is essential for growth of the rfaA mutant

A global study on the fitness of *B. subtilis* mutants ^18^ revealed that growth of the *rfaA* mutant is strongly impaired in minimal media in the absence of organic nitrogen sources. Addition of either glutamine or glutamate restored growth. A similar phenotype was also observed for mutants that are auxotrophic for glutamate due to the inability to synthesize this amino acid ^18,19^ (see Supplementary Fig. 1a). To verify the dependance of the *rfaA* mutant on glutamate, we cultivated the wild type strain *B. subtilis* 168 and the isogenic *rfaA* mutant GP1468 on C-Glc minimal medium in the absence and presence of glutamate. As shown in Fig. 1A, the wild type strain was able to grow on both media. In contrast, the *rfaA* mutant was not viable in the absence of glutamate. We also tested the effect of aspartate and glutamine. Whereas aspartate did not restore growth of the *rfaA* mutant, the presence of glutamine resulted in wild type-like growth of the *rfaA* mutant (see Supplementary Fig. 1b). Thus, glutamate and its direct precursor glutamine support growth of the mutant.

**Fig. 1.**
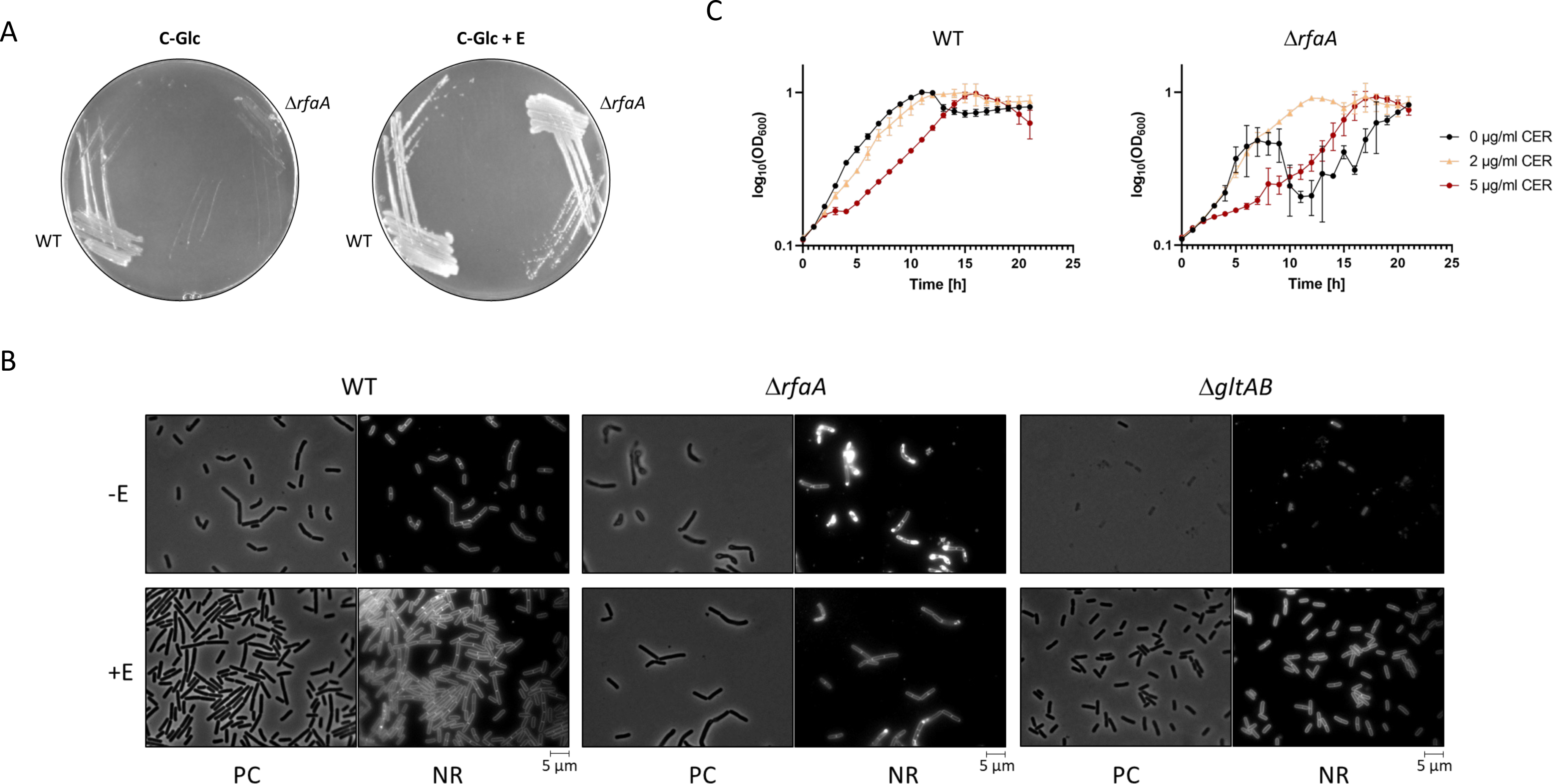
RfaA links fatty acid metabolism to glutamate availability. **a**, Growth assay of *B. subtilis* wild type and Δ*rfaA* mutant. Bacterial growth of the *B. subtilis* wild type strain 168 and the Δ*rfaA* mutant GP1468 has been assessed on minimal medium in absence or in presence of glutamate (10 mM), respectively. Bacterial growth was analyzed after incubation at 37 °C for 24 hours. **b**, Analysis of the cell morphology via microscopy. *B. subtilis* wild type strain (WT), and the mutant strains GP1468 (Δ*rfaA*) and GP807 (Δ*gltAB*) were precultivated in minimal medium supplemented with 10 mM glutamate and grown to an exponential growth phase. Cells were washed in minimal medium without glutamate, subsequently reinoculated in medium with and without glutamate (+E and –E). Samples were stained with nile red after 3 hours of growth and then analyzed by microscopy. **c**, Growth analysis of *B. subtilis* wild type and Δ*rfaA* mutant in presence of increasing cerulenin concentrations. Cells were inoculated to an OD_600_ of 0.1 and growth monitored in minimal medium in presence of 0 µg/ml (black), 2 µg/ml (yellow) and 5 µg/ml (red) cerulenin for 22 hours. Restricted growth of the Δ*rfaA* mutant in absence of cerulenin results from accumulation of toxic compounds. Abbreviations: E, glutamate; CER, cerulenin; PC, phase contrast; NR, nile red.

The observations presented above suggest that the *rfaA* mutant might be auxotrophic for glutamate. However, the biosynthetic pathway for glutamate in *B. subtilis* as well as its regulation have been extensively studied, and there is no indication that proteins in addition to those already identified might play a role in glutamate biosynthesis in *B. subtilis* ^19,20,21^. Therefore, we considered an alternative possibility, i. e. that growth in the absence of glutamate might result in a growth inhibition due to the accumulation of toxic compounds. To get a first impression on the role of glutamate for the *rfaA* mutant, we explored the cell morphology of the wild type strain 168 as well as the isogenic Δ*rfaA* (GP1468) and Δ*gltAB* (GP807) mutants. As shown in Fig. 1B, the presence of glutamate did not affect the cell morphology of the wild type strain. In constrast, the *rfaA* mutant cells were elongated both in the absence and presence of glutamate, and the cells seem to burst upon prolonged cultivation without glutamate. The glutamate-auxotrophic *gltAB* mutant failed to multiply in the absence of glutamate, but there were no effects on the cell morphology. These observations suggest that the inability of the *rfaA* mutant to grow without glutamate might result from the accumulation of toxic products rather than from a defect in glutamate biosynthesis.

Previously, RfaA has been implicated in the control of fatty acid homeostasis, and the accumulation of lipophilic clusters was observed in the *rfaA* mutant ^13^. We therefore wondered whether the *rfaA* mutant might accumulate an excess of lipids in the absence of glutamate. To answer this question, we stained the cells with the fluorescent membrane dye Nile Red ^22^. The cells of the wild type strain and the *gltAB* mutant exhibited a decent staining intensity both in the presence and absence of glutamate (Fig. 1B) In both strains, small punctuated structures were stained by Nile Red suggesting that lipids are stored in these strains. In contrast and in agreement with previous observations ^13^, the *rfaA* mutant cells formed larger lipophilic droplets at the cell poles. Most importantly, the *rfaA* mutant accumulated large amounts of lipids in the absence of glutamate as indicated by the excessive stain. Thus, the *rfaA* mutant seems to be unable to balance lipid metabolism with the availability of glutamate, the major donor of amino groups in cellular anabolism ^20,23^.

If the observed accumulation of lipids in the *rfaA* mutant in the absence of glutamate would be the growth-inhibiting problem, one would expect that the inhibition of lipid biosynthesis might restore growth of the mutant in the absence of glutamate. To address this idea, we made use of cerulenin, an inhibitor of fatty acid synthesis that binds β-ketoacyl-ACP synthases, such as FabF, FabHA and FabHB ^16,24,25^ (see Fig. 1C). In agreement with our earlier observations, growth of the *rfaA* mutant GP1468 in the absence of glutamate was severely compromised. As expected and in agreement with published data ^16^, the inhibition of fatty acid biosynthesis by increasing concentrations of cerulenin resulted in a growth inhibition for the wild type strain. In contrast, growth of the *rfaA* mutant was rescued in the presence of a low concentration of cerulenin (2 µg/ml), and even at 5 µg/ml, the growth was better as compared to the absence of the inhibitor. These observations demonstrate that indeed the accumulation of fatty acids in the absence of glutamate is toxic for the *rfaA* mutant.

Taken together, these results suggest that the excessive production of fatty acids is the growth-limiting factor for the *rfaA* mutant during growth in the absence of glutamate. Thus, RfaA may link fatty acid biosynthesis to the availability of the general amino group donor glutamate.

### RfaA controls fatty acid biosynthesis in B. subtilis

Taking into account the accumulation of lipids in the *rfaA* mutants, we hypothesized that RfaA might directly control fatty acid acquisition in *B. subtilis*. To address this question, we compared the concentration of fatty acids in the wild type strain 168 and the *rfaA* mutant GP1468. As shown in Fig. 2A, the total fatty acid content was increased in the *rfaA* mutant by about one third as compared to the wild type. In addition, we determined the relative profiles of the major fatty acids in *B. subtilis* (Fig. 2B). In agreement with previous studies ^16,26^, we found that branched-chain C15 and C17 fatty acids account for the majority of the fatty acid pool in the wild type strain, whereas palmitic acid (C16) corresponds to only about 10%. Of the branched fatty acids, the C15 species was predominant as compared to the C17 species (C17-C15 ratio of 0.53). In the *rfaA* mutant, the fraction of branched versus straight fatty acids was not severely affected, however, the average chain length of the branched amino acids was increased (C17-C15 ratio of 0.75). In conclusion, these results demonstrate, that RfaA indeed plays a role in the control of fatty acid biosynthesis. The increased fatty acid concentration in the absence of RfaA suggests that the RfaA protein limits fatty acid biosynthesis, likely by controlling the committed initial step, the biosynthesis of malonyl-CoA by ACCase that is typically subject to regulation ^27^.

**Fig. 2.**
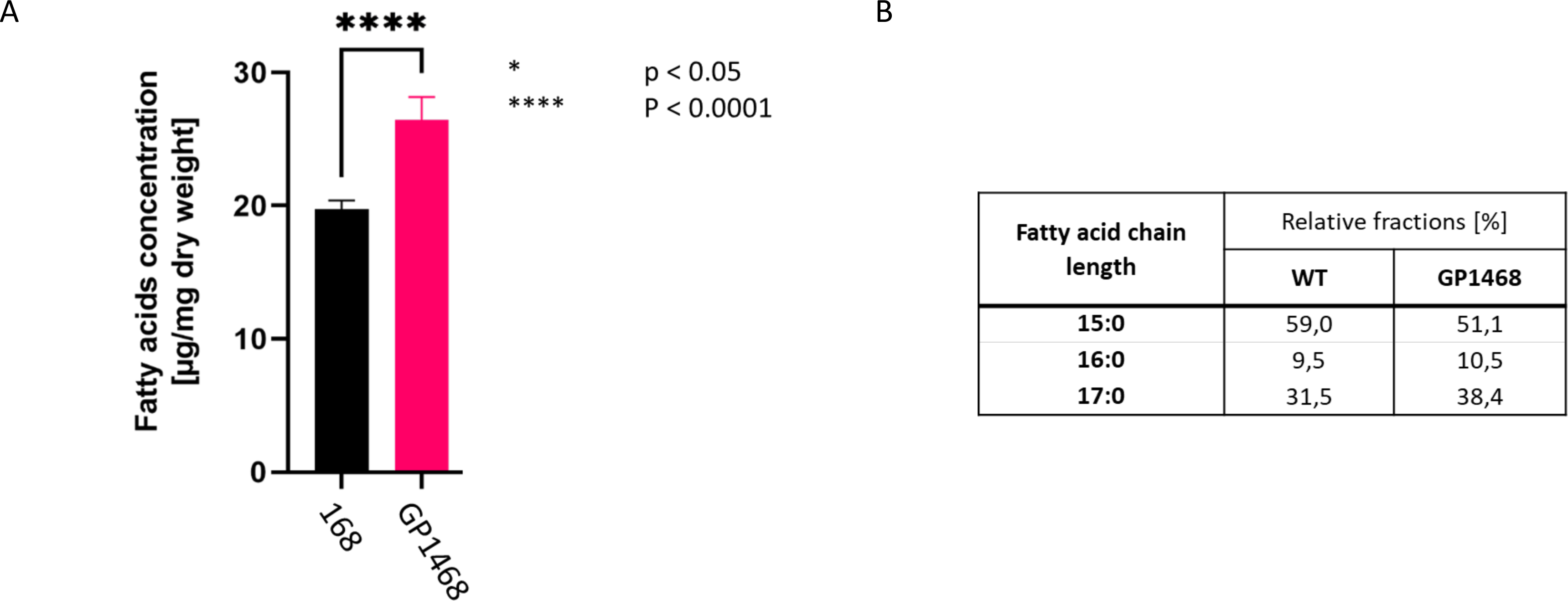
RfaA affects the cellular fatty acid content. **a**, Analysis of the total fatty acid content in the *B. subtilis* wild type strain 168, as well as the *rfaA* mutant GP1468 and the *rfaA* suppressor mutant GP3760. Cells were cultivated in complex medium and grown to an exponential phase. Fatty acid profiles were determined by GC-FID. One-way ANOVA tests were performed to calculate the p-values (p[168-GP1468]=0.00005; p[GP1468-GP3760]=0.0019). **b**, Relative fractions of the major fatty acids in *B. subtilis* were analyzed for the strains 168, GP1468 and GP3760. Similarly, data were obtained as described for the total fatty acid content shown in (**a**). Branched chain fatty acids C15 and C17 fatty acids include both iso and anteiso conformations. Abbreviation: %, percent.

### RfaA interacts with ACCase subunits

We assumed that RfaA might act by interaction with other proteins. Based on the conserved genomic clustering of the *rfaA* gene as a part of an operon with *accB* and *accC* encoding ACCase subunits (Fig. 3A) and its role in the regulation of fatty acid biosynthesis, the ACCase proteins were considered as prime candidates as interaction partners. Therefore, we used the bacterial two-hybrid system in which an adenylate cyclase is reconstituted if cloned proteins interact with each other resulting in β-galactosidase activity. As shown in Fig. 3B, both RfaA as well as AccB and AccC exhibited self-interaction. Moreover, we observed an interaction between the AccB and AccC subunits of ACCase. This observation is in excellent agreement the formation of an AccB_2_-AccC_2_ heterotetramer for the ACCase subunits ^28^. In addition, co-expression of RfaA and either AccB or AccC resulted in the reconstitution of a functional adenylate cyclase, thus confirming the interaction between the proteins. None of the three proteins showed an interaction with the Zip protein, which was used as the negative control. Thus, the RfaA-AccB and RfaA-AccC interactions are specific.

**Fig. 3.**
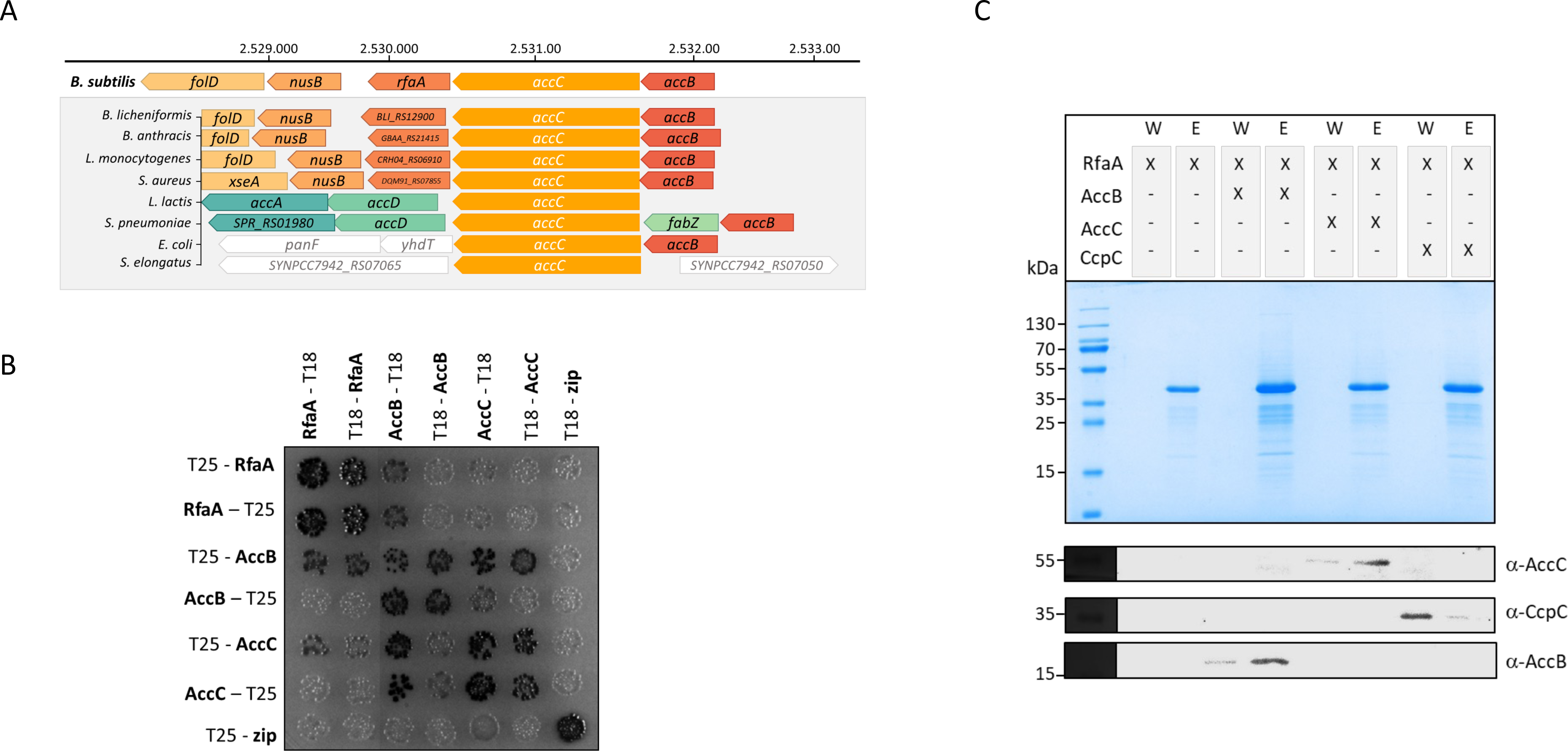
RfaA interacts with the ACCase subunits AccB and AccC. **a**, The genomic background of the *rfaA* gene in *B. subtilis* and the conservation of the genetic neighbourhood. The *rfaA* gene shares an operon with the *accB* and *accC* gene encoding two protein subunits of the ACCase. The analysis of the gene neighbourhood and conservation among different bacteria was done using the FlaGs tool ^63^. **b**, Bacterial two-hybrid (BACTH) assay to test for the interaction between RfaA and the ACCase proteins AccB and AccC. N-and C-terminal fusions of RfaA, AccB and AccC to the T18 or T25 domainn of the adenylate cyclase (CyaA) were created and the proteins were tested for interaction in *E. coli* BTH101. Dark colonies indicate an interaction that results in adenylate cyclase activity and subsequent expression of the reporter β-galactosidase. **c**, *In vitro* pulldown experiment with the GST-tagged RfaA. GST-RfaA was immobilized onto a glutathione sepharose column and incubated with purified AccB, AccC, or the control protein CcpC. The presence of GST-RfaA was analyzed by SDS PAGE, and the eluate and wash fractions were analyzed by western blot analysis using antibodies raised against AccB, AccC and CcpC, respectively. Abbreviations: GST, glutathione-S-transferase.

To further investigate the interaction between RfaA and the ACCase subunits, we assayed the binding of purified AccC and an *E. coli* cell extract containing overexpressed AccB (due to poor protein yields in AccB purification attempts) as well as the purified CcpC protein (as a negative control) to immobilized GST-tagged RfaA. As shown in Fig. 3C, both AccB and AccC co-eluted with RfaA whereas CcpC did not. These results confirm the specific interaction between RfaA and AccB and AccC and suggest that this interaction controls the activity of the ACCase subunits.

### RfaA controls the stability of AccC

The observed control of fatty acid synthesis by RfaA as well as the physical interaction between RfaA and the AccB and AccC proteins of the ACCase complex suggest that RfaA might control the activity of the ACCase. To test this idea, we purified the *B. subtilis* ACCase subunits, the RfaA protein, and the malonyl-CoA reductase to which the reaction was coupled. The AccB protein depends on its biotinylation for activity. To verify the biotinylation of the protein purified from *E. coli*, we assayed binding of the protein to streptavidin. This analysis indicated that AccB was present in the biotinylated form and could thus be used for activity assays. In the assay, the ACCase produced malonyl-CoA which was then determined in a coupled reaction with the malonyl-CoA reductase by assaying the oxidation of NADPH. As shown in Fig. S2, the ACCase was active; however, RfaA did not affect enzyme activity.

RfaA did not directly control ACCase actvity, thus suggesting a more indirect regulation of fatty acid synthesis by RfaA. Taking into account the physical interaction between RfaA and AccB and AccC, we considered the possibility that RfaA might control the stability of the ACCase subunits. To test this hypothesis, we analyzed the stability of AccB and AccC after growth in the absence and presence of glutamate. For this purpose, the strains were cultivated in minimal medium with and without glutamate, cell extracts were prepared ad aliquoted. To one aliquot of each extract, protease inhibitor was added. As shown in Fig. 4A, the levels of AccC in the absence of glutamate were affected by proteolysis in the wild type strain, as the signal for AccC was stronger in the presence of protease inhibitor. In contrast, no indications of proteolytic degradation of AccC were observed if the bacteria were grown in the presence of glutamate. AccB and the control protein GapA were not subject to proteolysis in the absence of glutamate. Thus, AccC seems to be subject to glutamate-dependent degradation.

**Fig. 4.**
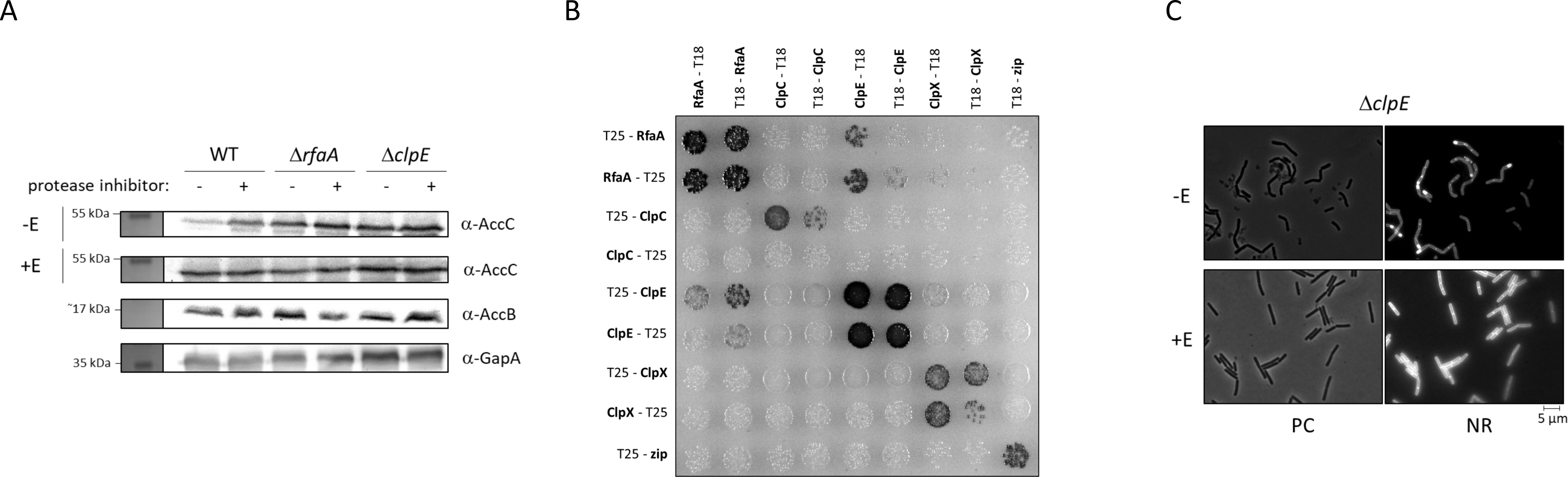
The AccC protein subunit of the ACCase is degraded in the absence of glutamate. **a**, Assay of the intracellular protein levels of the *B. subtilis* wild type (WT) and the *rfaA* mutant (GP1468) and the *clpE* mutant (GP3784). Cells were cultivated in presence of glutamate, then washed in minimal medium without glutamate and grown in medium with and without glutamate, respectively (-E, +E). Cultures were harvested after 3 h, cells were lyzed and the obtained protein extracts treated with and without protease inhibitor for 1 h, respectively. 10 µg of each samples were analyzed by Western blot analysis using antibodies raised against AccC (α-AccC), AccB (α-AccB) and GapA (α-GapA, negative control), respectively. **b**, Bacterial two-hybrid (BACTH) assay to test for the interaction between RfaA and the Clp proteins ClpC, ClpE and ClpX. N-and C-terminal fusions of RfaA, ClpC, ClpE and ClpX to the T18 or T25 domainn of the adenylate cyclase (CyaA) were created and the proteins were tested for interaction in *E. coli* BTH101. Dark colonies indicate an interaction that results in adenylate cyclase activity and subsequent expression of the reporter β-galactosidase. Abbreviations: E, glutamate. **c**, Analysis of the cell morphology via microscopy. *B. subtilis* mutant strain GP3784 (Δ*clpE*) was precultivated in minimal medium supplemented with 10 mM glutamate and grown to an exponential growth phase. Cells were washed in minimal medium without glutamate, subsequently reinoculated in medium with and without glutamate (+E and –E). Samples were stained with nile red after 3 hours of growth and then analyzed by microscopy. Abbreviations: E, glutamate, PC, phase contrast; NR, nile red.

Next, we asked whether the degradation of AccC in the absence of glutamate might be mediated by RfaA. For this purpose, we tested the AccC stability in the *rfaA* mutant GP1468. In contrast to the wild type strain, the amounts of AccC were similar, irrespective of the addition of a protease inhibitor (Fig. 4A), suggesting that RfaA is required for the glutamate-dependent degradation of AccC. The absence of AccC degradation in the *rfaA* mutant may be responsible for the observed accumulation of fatty acids in the mutant.

### RfaA modulates degradation of AccC by ClpEP

As described above, AccC is subject to RfaA-mediated proteolysis in the absence of glutamate. Condition-dependent proteolysis in bacteria is mainly catalyzed by complexes between the ClpP peptidase and an AAA+ protein unfoldase. In *B. subtilis*, ClpP has three AAA+ partner proteins, ClpC, ClpE, and ClpX ^29,30^. To test, which of these proteins might be involved in the RfaA-mediated degradation of AccC, we first analyzed the possible interactions between RfaA and the three AAA+ unfoldases. While all three proteins showed the well-established self-interaction, RfaA interacted exclusively with ClpE, suggesting that this protein might be the responsable ATPase subunit of the protease complex for AccC degradation (see Fig. 4B). To test this idea, we assayed the stability of AccC in the *clpE* mutant strain GP3784. As shown in Fig. 4A, AccC was similarly stable in the *rfaA* and *clpE* mutant strains, and not affected by the protease inhibitor. In conclusion, these observations suggest that the ClpEP protease complex degrades AccC in an RfaA-dependent manner in response to the availability of glutamate. If this hypothesis would be correct, one would assume that the *clpE* mutant would also accumulate lipids as observed for the *rfaA* mutant. Indeed, as in the *rfaA* mutant, we observed the formation of lipid droplets in the *clpE* mutant strain GP3784 (see Fig. 4C). Thus, RfaA and ClpE are both required for the degradation of AccC in the absence of glutamate, and the loss of either protein causes toxic lipid accumulation.

### Glutamate affects the protein-protein interaction between RfaA and ClpE

All results presented so far indicate that the RfaA-mediated degradation of AccC takes place only in the absence of glutamate but not if this amino acid is available in the medium. We therefore asked how this regulatory mechanism might respond to the availability of glutamate. The most obvious hypothesis suggests that glutamate might affect the interaction between RfaA and AccC and/or ClpE. To test this idea, we first coupled His-SUMO-tagged RfaA to a Ni^2+^-NTA column, and applied purified Strep-tagged AccB and AccC to the column in the presence or absence of glutamate. As shown in Fig. 5A, both proteins co-eluted with RfaA irrespective of the presence of glutamate. To test the interaction between ClpE and RfaA, we coupled His-tagged ClpE to a Ni^2+^-NTA column and then applied an *E. coli* cell extracts with overexpressed Strep-RfaA or Strep-RnpM. As shown in Fig. 5B, RfaA was retained on the column and co-eluted with ClpE in the absence of glutamate, whereas no co-elution was observed if glutamate was present. Moreover, the RNA-binding protein RnpM did not co-elute with ClpE in either condition, indicating the specificity of the assay. In conclusion, glutamate inhibits the interaction between RfaA and ClpE, thus preventing the degradation of AccC and allowing continued fatty acid biosynthesis to sustain growth. In contrast, AccC, RfaA, and ClpE interact in the absence of glutamate thus allowing the degradation of AccC and preventing the accumulation of fatty acids to toxic levels (see Fig. 6).

**Fig. 5.**
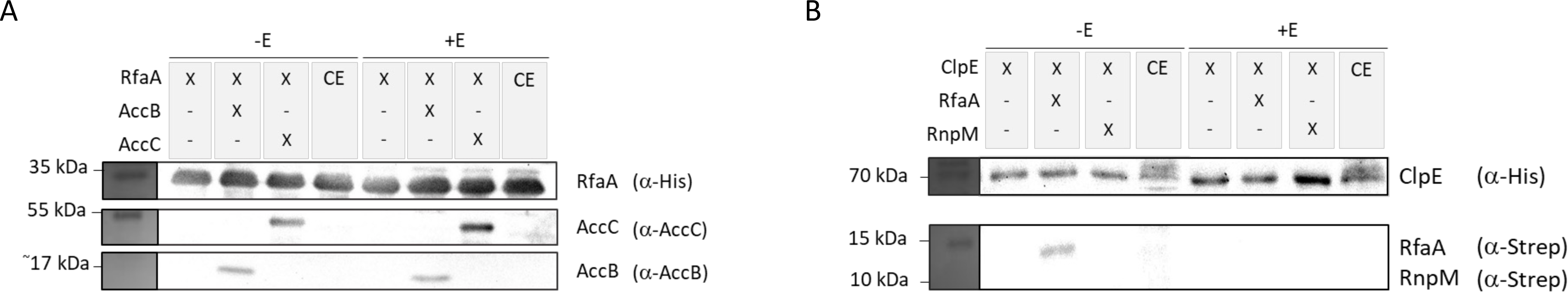
Effect of glutamate on the RfaA-AccBC and the RfaA-ClpE interactions. **a**, *In vitro* pulldown experiment with His-Sumo-tagged RfaA in presence and absence of glutamate. RfaA was immobilized onto a Ni-NTA agarose column and incubated with purified AccB and AccC with and without glutamate, respectively. The presence of His-Sumo-RfaA in the cell extract (CE) and the eluate fractions were analyzed by western blot analysis using antibodies raised against the His-tag, and the proteins AccB and AccC, respectively. **b**, *In vitro* pulldown experiment with His -tagged ClpE in presence and absence of glutamate. His-ClpE was immobilized onto a Ni-NTA agarose column and incubated with *E. coli* cell extract expressing Strep-RfaA or Strep-RnpM (negative control) in presence and absence of glutamate, respectively. The presence of His-ClpE in the cell extract (CE) and the presence of Strep-RfaA in the eluate fractions were analyzed by western blot analysis using antibodies raised against the His-tag and the Strep-tag, respectively. Abbreviations: E, glutamate; CE, cell extract.

**Fig. 6.**
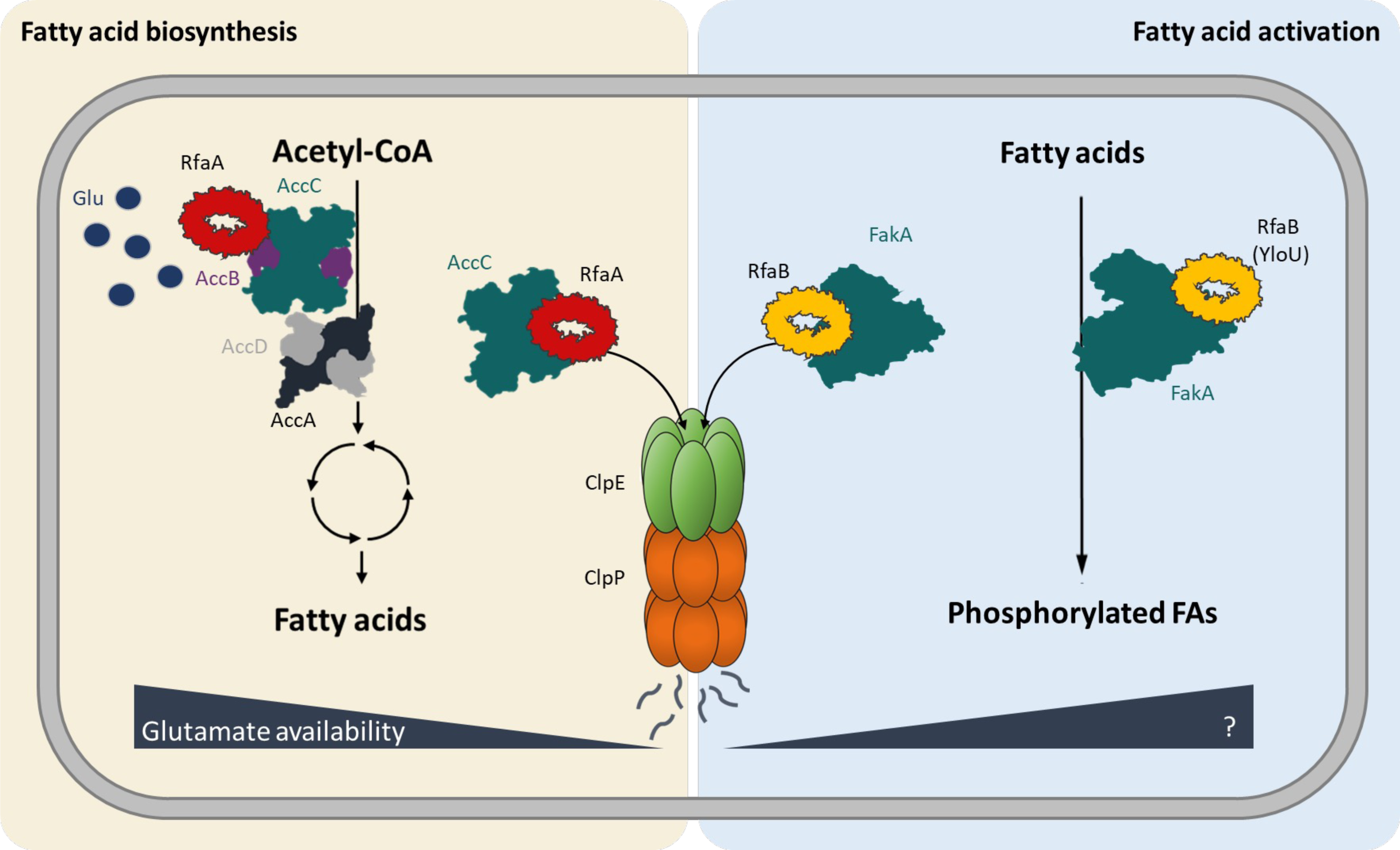
Proposed model of protease-mediated regulation of initiating steps of the fatty acid acquisition. The ACCase complex initiates the *de novo* fatty acid biosynthesis by catalyzing the conversion of acetyl-CoA to malonyl-CoA. In this work, the RfaA protein was identified as a novel regulator of this first committed step. In the absence of glutamate RfaA facilitates the ClpEP-dependent degradation of the AccC protein, thus, leading to an arrest of fatty acid biosynthesis. Based on that, a similar mechanism can be proposed for the regulation of the fatty acid activaton that is catalyzed by the fatty acid kinase FakA. The so far unknown paralogogous RfaB (YloU) may trigger the ClpEP-dependent degradation of FakA under conditions yet to be characterized. RfaA and RfaB are displayed as multimeric ring structures according to ColabFold predictions ^64^. To determine the oligomerization state, predicitions with highest scoring ipTM values were obtained for 10-, 11-and 12-meric RfaA proteins. Taking into consideration that ClpE exists as a hexameric ring in its active form, we considered the formation of dodecameric rings by RfaA and RfaB likely. Abbreviation: Glu, Glutamate.

## Discussion

In this work, we report a novel link between glutamate availability and lipid biosynthesis in the Gram-positive model organism *B. subtilis*. The observation that at low external glutamate concentrations, the biotin carboxylase subunit (AccC) of the acetyl-CoA carboxylase that catalyzes the committed step in fatty acid biosynthesis is subject to proteolytic degradation to protect the cell from toxic lipid accumulation shows how seemingly non-related branches of metabolism are intertwined. Together, our data allow to develop a model (Fig. 6) in which the previously unknown RfaA adaptor protein targets AccC for degradation to the ClpEP protease complex at low glutamate concentrations. In contrast, RfaA and AccC also interact at high glutamate concentration, but no interaction between RfaA and ClpE is possible under this condition thus allowing efficient fatty acid biosynthesis.

Glutamate is the most abundant metabolite in any living cell ^31^. In addition to its role as bulding block for proteins, gutamate is the central amino group donor for nearly all nitrogen-containing compounds including all amino acids. Moreover, this amino acid is derived from and can be degraded to oxaloacetate, an intermediate of the citric acid cycle. Thus, glutamate is the direkt metabolic link between catabolic and anabolic reactions as well as between carbon and nitrogen metabolism ^20,23^. Moreover, glutamate is the counterion for potassium, the most abundant metal ion in all cells ^32^. The direct link between glutamate availability and fatty acid biosynthesis is very important for the cell. If glutamate is available in addition to suitable carbon sources, the cells have all the resources to grow rapidly and thus also the need for an efficient lipid biosynthesis. In contrast, lipid accumulation in the absence of glutamate can be problematic for the cell if no counter-measures are taken (see Fig. 1).

The control of ACCase by protein-protein interactions has already been demonstrated in plant and algal chloroplasts and cyanobacteria as well as in proteobacteria such as *E. coli* ^33,34,35,36,37^. In all these organisms the biotin carboxyl carrier subunit (AccB) of the ACCase interacts with a widely conserved regulatory PII protein. PII proteins sense the energy state of the cell as well as the nitrogen supply by binding adenine nucleotides and 2-oxoglutarate ^38^. Binding of the PII proteins directly inhibits the biotin carboxyl carrier and thus ACCase activity. Under conditions of nitrogen limitation (perceived as accumulation of 2-oxoglutarate), the PII-2-oxoglutarate complex is unable to bind and inhibit ACCase ^38^. It has been suggested that this regulation serves to allow the formation and storage of fatty acids under conditions of nitrogen limitation. The PII proteins are conserved in archaea, many groups of bacteria, and plant chloroplasts. Interestingly, there is a negative correlation between the presence of PII proteins and proteins of the Asp23 family in bacteria. While the Asp23 proteins are present in most or all representatives of the chlamydiae, the Deinococcus-Thermus group, the firmicutes, fusobacteria and the thermotogae, the PII proteins are lacking or only present sporadically in these bacterial groups. On the other hand, a high prevalence of PII proteins correlates with absence or only partial presence in other groups such as the proteobacteria or the cyanobacteria ^10^. It is thus tempting to speculate that bacterial evolution has resulted in two principal mechanisms of ACCase regulation: control of ACCase activity by direct protein-protein interaction with PII in proteobacteria and cyanobacteria and control of ACCase degradation by RfaA-mediated targeting of the biotin carboxylase subunit to the Clp protease as described here for firmicutes.

The control of ACCase activity by PII has been studied *in vitro*; however, the regulatory mechanisms have not been investigated under physiological conditions. As mentioned above, in all studies Apo-PII inhibited ACCase activity whereas the addition of 2-oxoglutarate abolished the inhibition ^38^. Apo-PII is present in the cell if glutamine is available, and the binding of 2-oxoglutarate indicated a poor glutamine supply ^39^. Thus, PII seems to inhibit ACCase activity if glutamine is abundant whereas RfaA targets AccC for degradation if the nitrogen source is limiting (see Fig. 6). This obvious contradiction may reflect differences in nitrogen acquisition in these organisms. A difference between the organisms that use PII or RfaA to control malonyl-CoA biosynthesis might be in their mechanisms of nitrogen assimilation. The former bacteria (such as cyanobacteria and enterobacteria) have an anabolically active glutamate dehydrogenase for glutamate biosynthesis in addition to the conserved glutamine synthetase/glutamate synthase pathway ^40,41^. In contrast, *B. subtilis* can use only the latter pathway for the primary assimilation of ammonium ^19,42^. This difference in nitrogen acquistion may affect the use of lipids to store carbon under conditions of nitrogen limitation: the glutamate dehydrogenase reaction comes at no cost for the cell making it attractive to store extra carbon if nitrogen is abundant. In contrast, the glutamine synthetase reaction requires the hydrolysis of ATP, and this may disfavor lipid storage.

As mentioned above, most organisms that contain RfaA paralogs, have two or more of these proteins. In *B. subtilis*, these are RfaA and the YloU protein. Interestingly, YloU is encoded in a strongly conserved operon with FakA, the fatty acid kinase. This enzyme serves for the acquisition of fatty acids from the environment ^11,12^. A global analysis of protein stability in *B. subtilis* has revealed that FakA is degraded under conditions of glucose limitation, and that this degradation depends completely on the ClpP protease subunit whereas the ATPase subunits ClpC and ClpX had no effect ^43^. This suggests, that ClpE might also be the respnsible unfoldase for FakA degradation. It is tempting to speculate that in this case YloU serves as the adaptor protein to target FakA for ClpEP-dependent degradation. We therefore suggest renaming YloU to RfaB (regulator of fatty acid <u>a</u>cquisition B) (see Fig. 6).

So far, adaptor proteins have been identified that target proteins for degradation to the ClpCP and ClpXP complexes. In contrast, no adaptor protein for the ClpEP unfoldase-protease complex has been identified. Our work demonstrates that RfaA interacts with ClpE under conditions of glutamate limitation (Fig. 5B) and that this interaction is responsible for the degradation of AccC. Moreover, based on the degradation profile of FakA in the *clp* mutant strains ^43^, the paralogous RfaB (YloU) protein is likely to trigger ClpEP-dependent degradation of the fatty acid kinase FakA (see Fig. 6).

In many metabolic pathways, the enzymes that catalyze the committed step are subject to regulation. Our work underlines that protein degradation is an important mechanism to achieve this regulation. As described here for the AccC subunit of ACCase, the MurAA enzyme that catalyzes the committed step in peptidoglycan biosynthesis, is degraded by the ClpCP protease complex with ReoM serving as the adaptor protein ^43,44,45^. It is tempting to speculate that this mechanism is used to control more metabolic pathways than so far realized.

## Methods

### Strains, media and growth conditions

*E. coli* DH5α and Rosetta DE3 ^46^ were used for cloning and for the expression of recombinant proteins, respectively. All *B. subtilis* strains used in this study are derivatives of the laboratory strain 168. The strains are listed in Supplementary Table S1. *B. subtilis* and *E. coli* were grown in Luria-Bertani (LB) medium ^46^. For growth assays and the microscopy assay, *B. subtilis* was cultivated in C-Glc minimal medium ^19^. The media were supplemented with ampicillin (100 µg/ml), kanamycin (50 µg/ml), chloramphenicol (5 µg/ml), or erythromycin and lincomycin (2 and 25 µg/ml, respectively) if required.

### Phenotypic characterization

To assay growth of *B. subtilis*, the bacteria were inoculated in LB medium and precultured in C-Glc medium supplemented with 10 mM glutamate. The cultures were grown until exponential phase, harvested, washed three times in C-Glc medium without glutamate before an optical density at 600 nm (OD_600_) was adjusted to 1.0. For growth analysis on solid medium, the cells were streaked on C-Glc agar and C-Glc agar supplemented with either 10 mM of glutamate, 10 mM glutamine or 10 mM aspartate and incubated for 24 h at 37 °C. Similarly, the growth was assessed using a serial drop dilution assay. The dilution series was prepared using the same cells as described above and then spotted onto C-Glc agar plates with glutamate, glutamine and aspartate (final concentration 10 mM for all amino acids used), respectively. For growth analysis in liquid medium in presence or absence of cerulenin, the cells were precultured, harvested and washed as described above. The cells were adjusted to a final OD_600_ of 0.1, then used to inoculate a 96 well plate (Microtest Plate 96 Well, Sarstedt) containing C-Glc medium with the required cerulenin concentrations. Growth was tracked in an Epoch 2 Microplate Spectrophotometer (BioTek Instruments) at 37°C with linear shaking at 237 cpm (4 mm) for 22 h, and an OD_600_ was measured in 10 min intervals.

### Analysis of the cell morphology by microscopy

The *B. subtilis* strains were precultured, harvested and washed as described for the phenotypic characterization. The cells were then adjusted to an OD_600_ of 0.1 and re-inoculated in either C-Glc medium with glutamate (10 mM) or without glutamate and cultured at 37 °C. After 3 h, the cells were harvested and resuspended in 500 µl PBS pH 7.5. The cell suspension was supplemented with 25 µl of a 100 µg/ml nile red solution to stain bacterial membranes. 5 µl of the cells were then added to microscope slides coated with a thin layer of 1 % agarose and covered with a glass plate. Phase contrast pictures were obtained using the Axioskop 40 microscope equipped with an EC Plan-NEOFLUAR 100X/1.3 objective (Carl Zeiss, Göttingen, Germany) and the digital camera AxioCam MRm. Filterset 43 (EX BP 545/25, FT 579, EM BP 605/70; Carl Zeiss) was applied for nile red detection. The resulting images were processed using the AxioVision Rel 4.8 software.

### DNA manipulation

Transformation of *E. coli* and plasmid DNA extraction were performed using standard procedures ^46^. All commercially available plasmids, restriction enzymes, T4 DNA ligase and DNA polymerases were used as recommended by the manufacturers. Chromosomal DNA of *B. subtilis* was isolated as described ^47^. *B. subtilis* was transformed with plasmid and genomic DNA according to the two-step protocol ^47^.

### Construction of mutant strain by allelic replacement

Deletion of the *clpE* gene was achieved by transformation of *B. subtilis* 168 with a PCR product constructed using oligonucleotides to amplify DNA fragments flanking the target genes and an appropriate intervening resistance cassette as described previously ^48^. The integrity of the regions flanking the integrated resistance cassette was verified by sequencing PCR products of about 1,100 bp amplified from chromosomal DNA of the resulting mutant strain GP3784.

### Plasmid constructions

The *rfaA* allele was amplified using chromosomal DNA of *B. subtilis* 168 as the template and appropriate oligonucleotides that attached specific restriction sites to the fragment. Those were: BamHI and EcoRI for cloning *rfaA* in pGEX-6P-1 (Cytiva, Massachusetts, USA). The resulting plasmid was pGP3644. For the overexpression of His-Sumo-tagged RfaA, *rfaA* was amplified using chromosomal DNA of *B. subtilis* 168 as the template and appropriate nucleotides that attached BsaI and XhoI restriction sites to the fragments and cloned between the BsaI and XhoI sites of the expression vector pET-SUMO (Invitrogen, Germany). The resulting plasmid was pGP3630. For the overexpression of Strep-tagged AccB and AccC, the *accB* and *accC* alleles were amplified as described above using the appropriate nucleotides that attached KpnI and BamHI for cloning *accB* in pGP172 ^49^, and NdeI and BamHI for cloning *accC* in pGP574 ^50^. The resulting plasmids were pGP1025 and pGP1027 for AccB and AccC, respectively. For the expression and purification of Stre-tagged AccA and AccD in *B. subtilis*, *accA* and *accD* alleles were amplified using appropriate nucleotides that attached BamHI and SalI for cloning *accA* and *accD* in pGP382^51^. The resulting plasmids were pGP1723 and pGP1724 for AccA and AccD, resprectively. For the overexpression of His-tagged AccB and AccC, the *accB* and *accC* alleles were amplified as described above using the appropriate nucleotides that attached BamHI and PstI for cloning *accB* in pWH844 ^52^, and BamHI and SalI for cloning *accC* in pWH844. The resulting plasmids were pGP2649 and pGP3646 for AccB and AccC, respectively. To construct plasmids for Bacterial-two hybrid experiments, the *clpC, clpX, rfaA, accB, accC* and *clpE* allels were amplified as described above. The respective restriction sites XbaI and KpnI were used for cloning of the BACTH vectors pUT18, pUT18-C, pKT25 and p25-N ^53^. The resulting plasmids were pBP202-pBP205 for *clpC*, pBP206-pBP209 for *clpX*, pGP1470-pGP1473 for *rfaA,* pGP1736-pGP1739 for *accB*, pGP1740-pGP1743 for *accC* and pGP2157-pGP2160 for *clpE*. All plasmids are listed in Supplementary Table S2.

### Protein expression and purification

*E. coli* Rosetta(DE3) was transformed with the plasmids for pTRc99A-mcr ^54^, pGP1025, pGP1027, pGP1329 ^13^ or pGP3650 ^55^, for expression of Strep-tagged MCR, AccB, AccC, RfaA, and RnpM, respectively. For purification of 6xHis-tagged proteins, *E. coli* DH5α was transformed with the plasmids pGP706 ^56^, pGP2649, pGP2690 ^57^ and pGP3646 encoding 6xHis-CcpC, 6xHis-AccB, 6xHis-BirA and 6xHis-AccC, respectively. Expression of the recombinant proteins was induced by the addition of isopropyl 1-thio-β-D-galactopyranoside (final concentration, 1 mM) to exponentially growing cultures (OD_600_ of 0.8) of *E. coli* carrying the relevant plasmid. For the expression and purification of the ACCase subunits AccA and AccD, the *B. subtilis* wild type strain 168 was transformed with the plasmid pGP1723 (Strep-AccA) and pGP1724 (strep-AccD), respectively. Cultures were grown to a stationary phase (OD_600_ of 1.5) to express the proteins. His-tagged proteins were purified in 1 x ZAP buffer (50 mM Tris-HCl, 200 mM NaCl, pH 7.5), if not stated otherwise, and Strep-tagged proteins in buffer W (100 mM Tris-HCl, 150 mM NaCl, 1 mM Na_2_EDTA, pH 8.0). Cells were lysed by four passes at 18,000 p.s.i. through an HTU DIGI-F press (G. Heinemann, Germany). After lysis, the crude extract was centrifuged at 100,000 x g for 60 min and then passed over a Ni^2+^nitrilotriacetic acid column (IBA, Göttingen, Germany) for 6xHis-tagged proteins, or a StrepTactin column (IBA, Göttingen, Germany) for purification of Strep-tagged proteins. The protein was eluted with an imidazole gradient or D-desthiobiotin (2.5 mM), respectively. After elution, the fractions were tested for the desired protein using SDS-PAGE.

The protein samples were stored at −80°C until further use (but no longer than 3 days). The protein concentration was determined according to the method of Bradford ^58^ using the Bio-Rad dye binding assay and bovine serum albumin as the standard.

### In vitro analysis of protein-protein interactions

In order to verify the interaction between RfaA and AccB, as well as RfaA and AccC, *E. coli* Rosetta (DE3) was transformed with pGP3644 (GST-RfaA) and the protein was overexpressed as described above. The cells were lyzed by four passes at 18,000 p.s.i. through an HTU DIGI-F press (G. Heinemann, Germany) using GST binding buffer (140 mM NaCl_2_, 2.7 mM KCl, 10 mM Na_2_HPO_4_, 1.8 mM KH_2_PO_4_, pH 7.3). The crude extract of 500 ml cell culture was passed over a Glutathione Sepharose column (Cytiva, Massachusetts, USA) and washed with GST binding buffer until the wash fractions appeared clear (confirmation with Bradford assay). Glutathione Sepharose matrix with bound GST-RfaA were then co-incubated with 0.1 mg of the purified proteins Strep-AccB, His-AccC and His-CcpC, (negative control), as well as *B. subtilis* wild type cell extract for 2 h at 4 °C, respectively. Again, after extensive washing, GST-RfaA, together with its potential binding partners, was eluted from the column using GST elution buffer (50 mM Tris-HCl, 10 mM reduced glutathione, pH 8.0). The elution fractions from the eluates were analyzed by SDS PAGE followed by Western blot analysis using dedicated antibodies.

To study the effect of glutamate on the interaction between RfaA and the ACCase subunits AccB and AccC, as well as on the interaction between RfaA and ClpE, *E. coli* Rosetta (DE3) was transformed with pGP3630 (6xHis-SUMO-RfaA) and pGP3695 (6xHis-ClpE), respectively, and the proteins were overexpressed as described above. Cells were split and lysed by using either buffer A (50 mM Tris-HCl, 0.1 M KCl, 5 mM MgCl_2_, 3.5 mM ATP) or (II) buffer A +5 mM glutamate. Crude extracts were passed over a Ni^2+^nitrilotriacetic acid column (IBA, Göttingen, Germany) and washed with either buffer A or buffer A +5mM glutamate until the fractions appeared clear. The Ni^2+^-NTA matrix with bound His-Sumo-RfaA protein was then co-incubated with the purified Strep-tagged proteins AccB or AccC, respectively, for 2 h at 4 °C under constant rotation. For the His-ClpE pulldown, the His-ClpE coupled to the Ni^2+^-NTA matrix was co-incubated with the Strep-tagged proteins RfaA and RnpM (negative control), respectively, and also co-incubated for 2 h at 4 °C. Purification was continued by extensive washing of the column using buffer A with and without glutamate. His-Sumo-RfaA, together with potential binding partners, was eluted using an imidazole gradient. To verify the presence of the potential interaction partners, Western blot analysis was performed using antibodies raised against the His-and the Strep-tag, AccB and AccC, respectively, as described ^59^.

### Comparison of protein levels by quantative Western blot analysis

To investigate potential proteolysis of the ACCase subunits, the protein levels were determined using a protease inhibitor assay. To assay a potential effect of glutamate on the proteolysis of AccB or AccC proteins, the *B. subtilis* wild type strain 168 was precultured in C-Glc medium and C-Glc medium supplemented with 10 mM glutamate. The cells were then adjusted to an OD_600_ of 0.1 and inoculated in either medium with or without glutamate and cultured at 37 °C until an OD_600_ of 0.8. Cells were harvested by centrifugation, and resuspended in Z buffer (60 mM Na_2_HPO_4_, 40 mM NaH_2_PO_4_, 10 mM KCl, 1 mM MgSO_4_, 50 mM β-Mercaptoethanol) supplemented with LD-mix (1% Lysozyme, 0.1% DNase I). To lyze the cells, the suspensions were incubated for 30 min at 37 °C with constant shaking followed by centrifugation at 14.000 rpm at 4 °C for 10 min. Protein concentration of the resulting cell extract was determined using Bradford assay. For the protease inhibitor assay, each cell extract (from cultures with and without glutamate, respectively) was split in two samples with similar protein concentration of which one was mixed with Complete Protease Inhibitor (Roche, Mannheim, Germany) and the other one with ddH_2_O as a control. The samples were incubated for 30 min at 37 °C. Protein levels of AccB and AccC were quantified using 10 µg total protein concentration of the samples for Western blot analysis with antibodies raised against the AccB and AccC protein, respectively.

To assay an effect of different mutant strains on the ACCase subunit protein levels, *B. subtilis* strains 168 (WT), GP1468 (Δ*rfaA*), GP3784 (Δ*clpE*) were precultured, harvested and washed as described for the phenotypic characterization. The cells were then adjusted to an OD_600_ of 0.1 and re-inoculated in C-Glc medium without glutamate and cultured at 37 °C. After 3 h, the cells were harvested and lyzed with Z buffer and LD-mix as described above. Similarly, the protein levels were quantified using Western blot analysis.

### Bacterial two-hybrid assay

Primary protein-protein interactions were identified by bacterial two-hybrid (BACTH) analysis ^53^. The BACTH system is based on the interaction-mediated reconstruction of *Bordetella pertussis* adenylate cyclase (CyaA) activity in *E. coli* BTH101. Functional complementation between two fragments (T18 and T25) of CyaA as a consequence of the interaction between bait and prey molecules results in the synthesis of cAMP, which is monitored by measuring the β-galactosidase activity of the cAMP-CAP-dependent promoter of the *E. coli lac* operon. Plasmids pUT18C and p25N allow the expression of proteins fused to the T18 and T25 fragments of CyaA, respectively. For these experiments, we used the plasmids pGP1470-pGP1473, which encode N-and C-terminal fusions of T18 or T25 to *rfaA*, pGP1736-pGP1739 for *accB*, pGP1740-pGP1743 for *accC*, pBP202-pBP205 for *clpC,* pGP2157-pGP2160 for *clpE* and pBP206-pBP209 for *clpX.* These plasmids were used for co-transformation of *E. coli* BTH101 and the protein-protein interactions were then analyzed by plating the cells on LB plates containing 100 µg/ml ampicillin, 50 µg/ml kanamycin, 40 µg/ml X-Gal (5-bromo-4-chloro-3-indolyl-ß-D-galactopyranoside), and 0.5 mM IPTG (isopropyl-ß-D-thiogalactopyranoside). The plates were incubated for a maximum of 36 h at 28°C.

### In vitro ACCase activity assay

To study the effect of RfaA on the activity of the ACCase complex, the conversion of acetyl-CoA to malonyl-CoA was assayed. This conversion was coupled to the reaction of the malonyl-CoA reductase (MCR) of *Chloroflexus aurantiacus* ^54^. ACCase protein components AccBC and AccDA were purified as described above. To ensure a catalytically active ACCase, biotinylation of AccB is required. Purified AccB protein was used for *in vitro* biotinylation. A reaction mix containing AccB (1 mg), *E. coli* cell extract expressing BirA (10 mg), and the reaction buffer (50 mM Tris-HCl, 100 mM KCl, 10 mM ATP and 1 mM biotin) was prepared and incubated for 1 h at 37 °C. The biotinylated AccB was purified again using a heparin column and the biotinylation was confirmed by Western blot analysis using Streptavidin-horseradish peroxidase.

The MCR uses malonyl-CoA as a substrate to produce 3-hydroxypropionate in a NADPH-dependent reaction ^60^. The oxidation of NADPH was monitored at 365 nm in a multi-well plate reader. The reaction was performed in a total volume of 150 µl containing the reaction buffer (50 mM Tris, 100 mM KCl, 20 mM MgCl_2_, 10 mM ATP, 0.5 mM DTT, 10 mM NaHCO_3_, 1 mM NADPH and 1.6 mM Acetyl-CoA), the purified MCR (5µg) and the purified proteins AccB, AccC and AccDA (100 nM each) as well as RfaA (2 µM). The reaction was initiated by addition of the MCR and monitored for 12 h at 25 °C. To verify the activity of the MCR, a set-up without the components of the ACCase and the substrate acetyl-CoA was prepared. This reaction was initiated by the addition of 0.3 mM malonyl-CoA and monitored similarly.

### Preparation of samples for fatty acid profiling

The *B. subtilis* wild type 168, *rfaA* mutant GP1468 and the *rfaA* suppressor GP3760 were inoculated in LB medium and grown over night at 37 °C. The freshly grown over night cultures were used to inoculate 200 ml LB medium cultures to an OD_600_ of 0.1. The cultures were grown at 37 °C to a final OD_600_ of 0.8, harvested by centrifugation at 5.000 rpm for 6 min and washed three times in PBS pH 7.5. The cell pellets were finally resuspended in PBS pH 7.5, snap frozen in liquid nitrogen and subjected to lyophilisation. The freeze-dried cells were then used to determine the fatty acid content. The preparation of methyl esters of fatty acids (FAMEs) was performed as described ^61^. For this purpose, 1 ml methanol:toluene:sulfuric acid:dimethoxypropane, 33:17:1.4:1, *v*/*v*/*v*/*v* with 50 µg tripentadecanoin as internal standard for later quantification was added to 5 mg freeze-dried cells. After 1 h incubation at 80°C, 1.5 mL saturated NaCl solution and 1.2 mL hexane were added subsequently. The hexane phase was dried under streaming nitrogen and dissolved in 10 µl acetonitrile. The gas chromatographic analysis with flame-ionization detection (GC-FID) was performed as described ^62^.

## Acknowledgements

We wish to thank Ostap Krynytskyy and Patrick Hitschrich for the help with some experiments. Anika Klewing and Jan Gundlach are acknowledged for the construction of some plasmids. This work was supported by the Deutsche Forschungsgemeinschaft (DFG) via SFB 1565 (Projektnummer 469281184 (P11 to J.S.).

## Contributions

D.W., I.F., and J.S. conceptualized the study. D.W., C.H., S.L., D.T., L.S., and F.M.C.. developed the methodology, performed the experiments and analyzed the data. D.W. and J.S. wrote the original draft of the manuscript. I.F., C.H. and F.M.C. reviewed and edited the manuscript. J.S. acquired funding. I.F., F.M.C., and J.S. provided supervision.

