## Supplementary Fig. for "RfaA (YqhY), a novel adaptor protein, controls metabolite-sensitive protein degradation in *Bacillus subtilis*"

S1

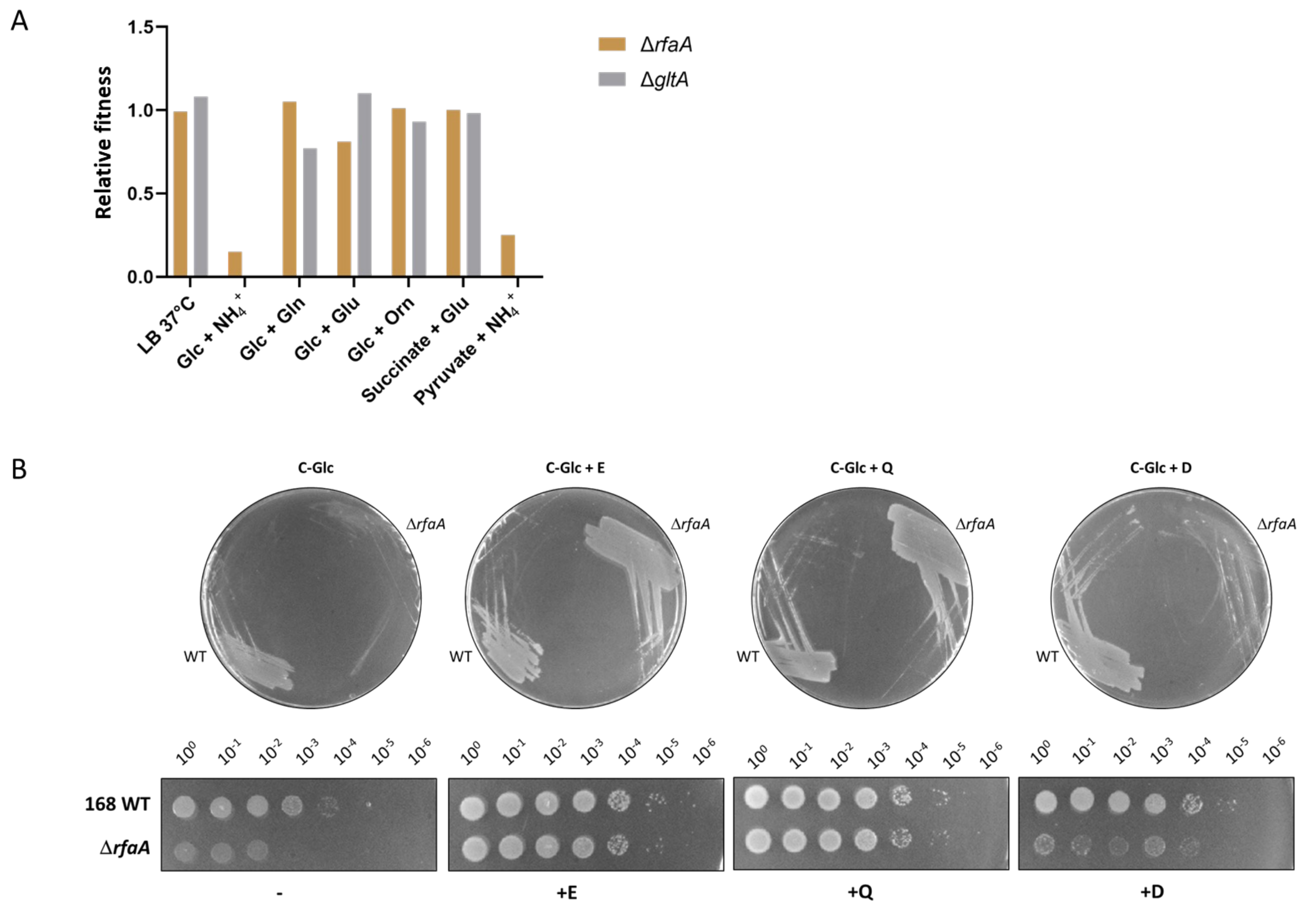


**Fig. S1. Glutamate is required for growth of the *B. subtilis* Δ*rfaA* mutant. a**, Comparison of the relative fitness of the Δ*rfaA and the* Δ*gltA* mutants in *B. subtilis*. Data used to compare the two strains was obtained from KOO. **b**, Growth assay of *B. subtilis* wild type and Δ*rfaA* mutant. Bacterial growth of the *B. subtilis* wild type strain 168 and the Δ*rfaA* mutant GP1468 has been assessed on minimal medium in absence or in presence of glutamate (E), glutamine (Q) and aspartate (D), respectively. Additionally the growth has been analyzed by drop dilution assays displayed below.

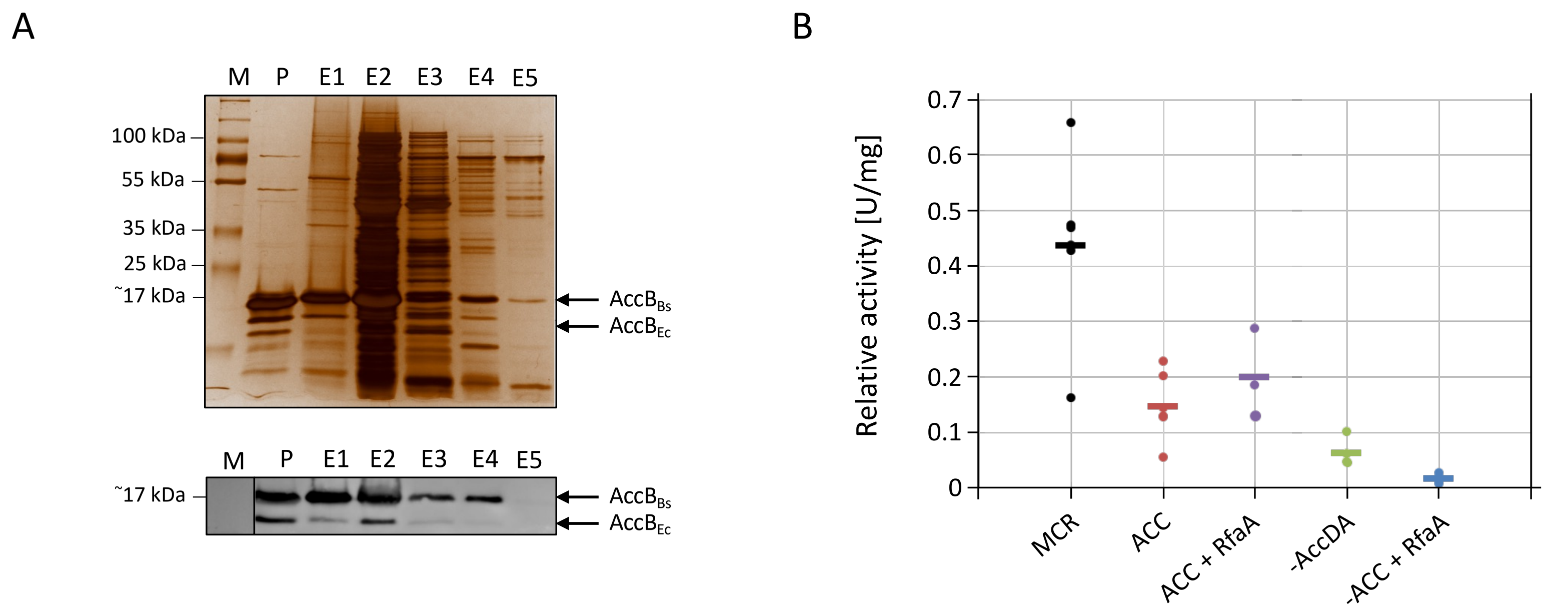


**Fig. S2. RfaA does not affect ACCase activity. a**, Confirmation of the biotinylation of AccB. The purified AccB protein (P) was used for *in vitro* biotinylation and purification via a heparin column. The presence of AccB was confirmed by SDS PAGE of the elution fractions (E1-E5) and subsequent silver staining. The biotinylation state has been analyzed by Western blot analysis using streptavidin-HRP. **b**, Activity of the ACCase complex was assayed by a coupled enzyme assay. The production of malonyl-CoA by the ACCase was coupled to the NADPH-dependend reaction of the malonyl-CoA reducatase (MCR). The oxidation of NADPH was monitored using a multi-well PlateReader. Dots represent single measurements and bars indicate the mean value of four replicates.

**Supplementary table S1. Strains used in this study.**

| Name | Genoytpe | Source or Reference |
|---|---|---|
| 168 | *trpC2* | Laboratory collection |
| GP807 | *trpC2* Δ*gltAB*::*tet* | 21 |
| GP1468 | *trpC2* Δ*rfaA*::*ermC* | 13 |
| GP3784 | *trpC2* Δ*clpE*::*neo* | This work |

**Supplementary table S2. Plasmids used in this study.**

| Plasmid | Relevant characteristics | Source or Reference |
|---|---|---|
| p25-N | N-term. fusion of the target protein to the T25 fragment of adenylate cyclase | Karimova |
| pBP202 | pUT18 with *clpC* | This work |
| pBP203 | pUT18C with *clpC* | This work |
| pBP204 | pKT25 with *clpC* | This work |
| pBP205 | p25-N with *clpC* | This work |
| pBP206 | pUT18 with *clpX* | This work |
| pBP207 | pUT18C with *clpX* | This work |
| pBP208 | p25-N with *clpX* | This work |
| pBP209 | pKT25 with *clpX* | This work |
| pGEX-6P-1 | Expression of N-term. GST-tagged proteins in *E. coli* | Cytiva, USA |
| pGP172 | Expression of N-term. Strep-tagged proteins in *E. coli* | 49 |
| pGP382 | Expression of C-term. Strep-tagged proteins in *B. subtilis* | 51 |

| | | |
|---|---|---|
| pGP574 | Expression of C-term. Strep-tagged proteins in *E. coli* | 50 |
| pGP706 | Expression of N-term. His-tagged CcpC | 56 |
| pGP1025 | Expression of N-term. Strep-tagged AccC | This work |
| pGP1027 | Expression of C-term. Strep-tagged AccB | This work |
| pGP1329 | Expression of N-term. Strep-tagged RfaA | This work |
| pPG1470 | pUT18 with *rfaA* | This work |
| pPG1471 | pUT18C with *rfaA* | This work |
| pPG1472 | p25-N with *rfaA* | This work |
| pPG1473 | pKT25 with *rfaA* | This work |
| pGP1723 | Expression of C-term. Strep tagged AccA in *B. subtilis* | This work |
| pGP1724 | Expression of C-term. Strep tagged AccD in *B. subtilis* | This work |
| pGP1736 | pUT18 with *accB* | This work |
| pGP1737 | pUT18C with *accB* | This work |
| pGP1738 | p25-N with *accB* | This work |
| pGP1739 | pKT25 with *accB* | This work |
| pGP1740 | pUT18 with *accC* | This work |
| pGP1741 | pUT18C with *accC* | This work |
| pGP1742 | p25-N with *accC* | This work |
| pGP1743 | pKT25 with *accC* | This work |
| pGP2157 | pUT18 with *clpE* | This work |
| pGP2158 | pUT18-C with *clpE* | This work |
| pGP2159 | p25-N with *clpE* | This work |
| pGP2160 | pKT25 with *clpE* | This work |
| pGP2649 | Expression of N-term. His-tagged AccB | This work |
| pGP3644 | pGEX-6P-1 with *rfaA* | This work |

| | | |
|---|---|---|
| pGP3646 | Expression of N-term. His-tagged AccC | This work |
| pGP3650 | Expression of N-term. Strep-tagged RnpM | 55 |
| pKT25 | C-term. fusion of the target protein to the T25 fragment of adenylate cyclase | 53 |
| pKT25-zip | C-term. fusion of the leucine zipper to the T25 fragment of adenylate cyclase (control) | 53 |
| pTrc99A-mcr | Expression of N-term. Strep-tagged MCR from *Chloroflexus aurantiacus* | 54 |
| pUT18 | N-term. fusion of the target protein to the T18 fragment of adenylate cyclase | 53 |
| pUT18-C | C-term. fusion of the target protein to the T18 fragment of adenylate cyclase | 53 |
| pUT18-zip | N-term. fusion of the leucine zipper to the T25 fragment of adenylate cyclase (control) | 53 |
| pWH844 | Expression of N-term. His-tagged proteins in *E. coli* | 52 |
